# A Computational Model for Storing Memories in the Synaptic Structures of the Brain

**DOI:** 10.1101/2022.10.21.513291

**Authors:** Vivek George, Vikash Morar, Gabriel Silva

## Abstract

Spike-timing dependent plasticity (STDP) is widely accepted as a mechanism through which the brain can learn information from different stimuli(1, 2). Basing synaptic changes on the timing between presynaptic and postsynaptic spikes enhances contributing edges within a network(3, 4). While STDP rules control the evolution of networks, most research focuses on spiking rates or specific activation paths when evaluating learned information(5–7). However, since STDP augments structural weights, synapses may also contain embedded information. While imaging studies demonstrate physical changes to synapses due to STDP, these changes have not been interrogated based on their embedding capacity of a stimulus(8–12). Here, we show that networks with biological features and STDP rules can embed information on their stimulus into their synaptic weights. We use a k-nearest neighbor algorithm on the synaptic weights of thousands of independent networks to identify their stimulus with high accuracy based on local neighborhoods, demonstrating that the network structure can store stimulus information. While spike rates and timings remain useful, structural embeddings represent a new way to integrate information within a biological network. Our results demonstrate that there may be value in observing these changes directly. Beyond computational applications for monitoring these structural changes, this analysis may also inform investigation into neuroscience. Research is underway on the potential of astrocytes to integrate synapses in the brain and communicate that information elsewhere(13–15). In addition, observations of these synaptic embeddings may lead to novel therapies for memory disorders that are difficult to explain with current paradigms, such as transient epileptic amnesia.

**Significance Statement:** Learning in the brain is often achieved via spike-timing dependent plasticity changing the structure of synapses to augment the strength between neurons. Typically, these changes contribute to other behaviors in the network, such as spiking rates or spike timings. However, observing these changes themselves may be fruitful for interrogating the learning capability of networks in the brain. Using a computational model, we demonstrate that the synaptic weights contain an embedding of the stimulus after a certain amount of recurrent activity occurs. It is possible that networks in the brain embed information in a similar way and that external readers, such as astrocytes, can interrogate, integrate, and transport this synaptic weight information to process stimuli.

---

Biological neural networks adapt to stimuli to learn through synaptic plasticity(1, 2). Whether through differences in firing rate, postsynaptic potential magnitude, or myelination, the brain requires state changes to transfer information(5– 7, 16–18). Spike-timing dependent plasticity (STDP) rules are crucial to these state changes, as they incorporate the activity of the network into the changes that occur. In both excitatory and inhibitory neurons, potentiation and depression are essential to adapting networks based on the stimulus presented to them(3, 4). However, excitatory and inhibitory neurons need different learning rules for potentiation and depression, as their synapses strengthen from opposite postsynaptic neuron outcomes(19). An excitatory neuron would experience synaptic potentiation if the downstream neuron fires soon after it did while an inhibitory neuron would experience synaptic potentiation if the downstream neuron never fires.

Both potentiation and depression are intertwined with the postsynaptic neuron’s refractory period. Through this, the refractory period plays a role in the development of structural changes within a network(20). The refractory period, which is the time after a neuron fires when it is incapable of firing again, temporarily prevents incoming signals from strengthening through potentiation. Indeed, when a signal from a presynaptic neuron reaches a refractory postsynaptic neuron, the synaptic weight between them is reduced. Fine-tuning of refractory periods is necessary to avoid over-reduction of weights while also ensuring that depression occurs when needed(21). Experimentally, it is challenging to change the refractory periods of real neurons to make observations about the impact of refraction. Computational models, however, can freely augment neuron refractory periods and observe how activity and activity-based learning rules are affected.

Over time, potentiation has been canonically credited for memory formation and learning in the brain. However, recent evidence has shown that depression can be important for learning as well, particularly for tasks involving object recognition(22–27). While depression is often used to stabilize weights after potentiation, depression can also be taken advantage of to reduce weights intentionally to embed information from a stimulus(28, 29). Though scientists can deactivate either potentiation or depression(30), there are few experiments interrogating how well learning takes place with just one or the other(31, 32). Without the limitations and external variables of *in vivo* and *in vitro* experiments, our model can further explore how well information can be embedded with just potentiation or depression.

When studying STDP, researchers generally observe neuron activity within biological networks. Therefore, much of the experimental work on STDP has been done to observe changes to the firing rates, connectivities, or postsynaptic potential magnitudes of neurons(3, 33). However, recent experiments show that in the process of causing changes to dynamics, STDP may also embed the stimulus into the morphological structure of the network itself(34, 35). As potentiation and depression result from activations caused by the stimulus, the resulting synaptic weight changes could reflect the stimulus. While limited by the neuroanatomical connectivity, potentiation and depression are appealing candidates for memory formation through changes at the dendrite(8–12). Though scientists can image dendritic spines as they grow and shrink from STDP(36, 37), a computational model can more easily quantify how the synapse is changing due to a stimulus. With this quantification, we can elucidate the effectiveness of the structural embedding created via STDP.

While potentiation and depression are effective for learning, they are limited by the number of active neurons inside the network. In particular, previous works have shown that an assembly of neurons has to be sufficiently large to reconstitute the stimulus presented to it(38, 39). Furthermore, one mechanism by which working memory can be trained is by increasing the number of neurons that respond to a given stimuli(40). Through these experiments, structural changes seem to be capable of embedding information and bringing a sufficient number of neurons together for learning new things.

In this work, we use the leaky integrate-and-fire neuron model(41) with refractory periods(42) and axonal delays to incorporate geometry into the network(43). These axonal delays were based on measurements of murine hippocampal axon lengths and conduction velocities(44, 45). In addition, we implemented inhibition within the network at a percentage also observed in the rat hippocampus(46). To modify the network based on the activity resulting from the axonal delays, we incorporated STDP rules for both excitatory edges(47) and inhibitory edges(4) based on murine hippocampal data.

Using this neuron model in a recurrent network, we demonstrate that the synaptic weights of the network contain significant information about the stimulus. In particular, since we designed our network to obey canonical neurobiological signaling principles, it follows that our model and a biological network of neurons embed stimulus information in a similar way.

Furthermore, the refractory periods within the model can be adjusted in ways that biological systems cannot be to observe the effects on learning. Moreover, we can isolate potentiation and depression to determine how well they can embed the stimulus independently as well as together. As the stimulus causes activations that modify synaptic weights, a concatenated vector of the weights evolves in time and reflects the stimulus in a particular time window. Based on the STDP weight changes, the trajectory of this vector changes within the state space of synaptic weights. This vector is our primary method of accurately quantifying the structural changes STDP implements over time. In this paper, we hypothesize that due to the STDP changes, this weight vector contains information about the stimulus that an external reader can access. Other works suggest that astrocytes may be able to integrate plasticity changes across thousands of synapses to read out information from neurons within the hippocampus(13, 48).

## 1. Methods

The overview of our methodology is highlighted in Figure 1.

**Fig. 1.**
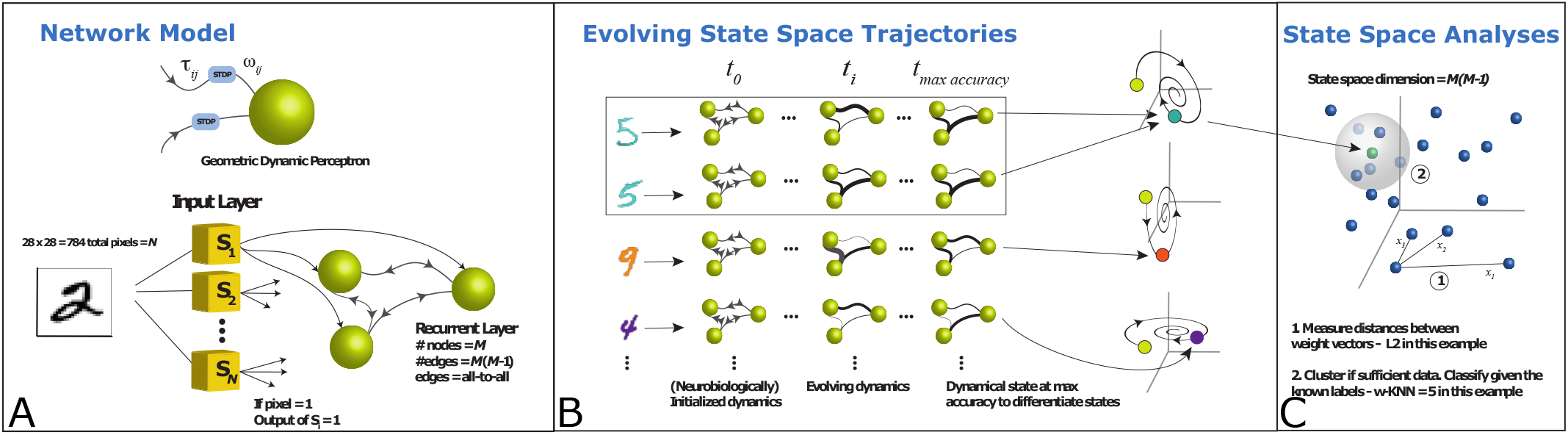
An overview of our network model and analysis methods. (A) The neuron model we employ in our network is a leaky integrate-and-fire with a time delay (length) and weight for each edge. MNIST images consisting of 784 pixels feed into the input layer with each input neuron corresponding to one pixel. Each input layer neuron connects to each recurrent layer neuron with a different delay. As the recurrent layer neurons activate, they send signals to each other with varying delays as well to propagate signals within the layer. As signals travel within the recurrent layer, a biologically-inspired STDP rule modulates the weights of each edge. (B) For our simulations, we initialize many identical networks before stimulating them with different MNIST images to run in parallel. As the simulations progress, the STDP rules result in similar modifications between networks stimulated by similar images. The weight values of each edge are concatenated into a weight vector and we notice that weight vectors from similar stimuli travel along similar trajectories. (C) During our simulation, we run these concatenated weight vectors through a k-nearest neighbors algorithm. We classify vectors from simulations with unknown stimuli based on their five nearest neighbors with known stimuli.

### A. Neuron Model

For this analysis method, we used an extension of the leaky integrate-and-fire neuron model to incorporate axon distances(41, 43). Each neuron accumulates membrane potential from arriving signals while its overall potential decays away (leaks) as a function of time. With the subscript *i* representing the postsynaptic neuron and subscript *j* representing any presynaptic neuron that has sent a signal, the membrane potential can be expressed as:

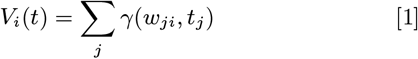

The membrane potential *V*_*i*_ equals the sum of the weights of each incoming signal *w*_*ji*_ after they run through the decay function *γ* based on their respective arrival times *t*_*j*_. Once the threshold potential is reached, the neuron fires along all its edges and becomes refractory with its membrane potential resetting(42).

Over time, the weights of each edge are modulated using spike-timing dependent plasticity (STDP) rules found in biological neurons(49). As excitatory and inhibitory edges need to be updated differently in response to network activity, a different STDP rule was used for each, as seen in Figure 2(4, 47). For excitatory edges, potentiation occurs when a postsynaptic neuron fires soon after a signal arrives and depression occurs when the postsynaptic neuron is refractory when a signal arrives. For inhibitory edges, potentiation (making the edge more inhibitory) occurs when a postsynaptic neuron does not fire soon after a signal arrives and depression occurs when a postsynaptic neuron fires near the signal arrival.

**Fig. 2.**
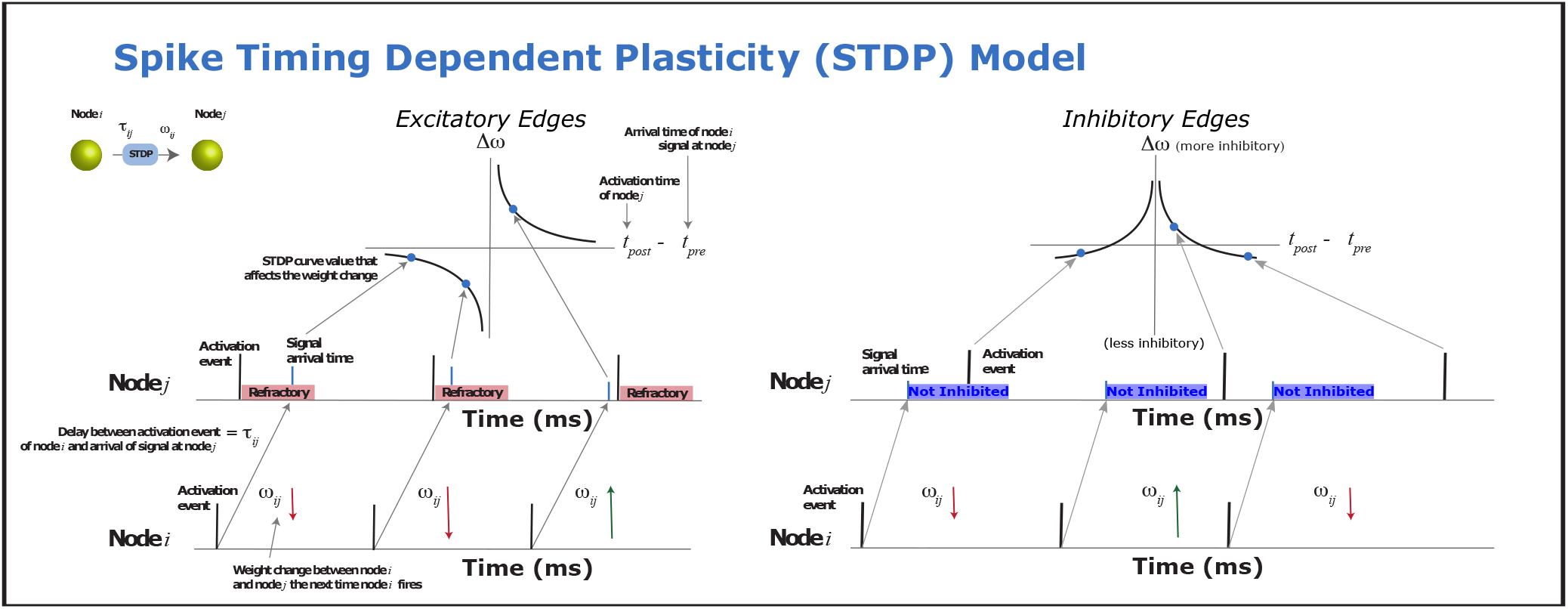
Here are the biologically-inspired STDP rules for the edges in our network. If an excitatory signal from a presynaptic neuron arrives at a postsynaptic neuron before it fires, the weight of the edge between them increases. If the signal arrives at the postsynaptic neuron while it is refractory, the weight of that edge decreases instead. Inhibitory edges are modified with a different biologically-inspired STDP rule. If the postsynaptic neuron fires close to when the presynaptic signal arrives, the weight of the inhibitory edge decreases. If a sufficiently long time passes after the presynaptic signal arrives without a postsynaptic activation, then the magnitude of the inhibitory edge increases.

In addition to arrival and departure times of signals, there are also parameters for the STDP change amplitudes and the time constants for scaling that time difference. These provide direct control over how large each weight change can be and how dependent each one is on the time difference observed.

For excitatory edges, all of this can be expressed as such:

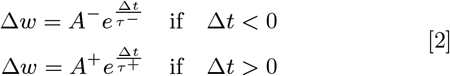

In this equation, Δt is the time of postsynaptic neuron firing minus the time of presynaptic neuron firing. *A*^+^ and *A*^−^ represent the amplitudes for potentiation and depression, respectively. *τ* ^+^ and *τ* ^−^ are the STDP time constants as previously described. Δw represents how much the edge weight will change due to that pair of firings as a result of STDP.

For inhibitory edges, weight changes can be expressed as such:

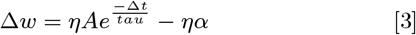

Similar to the excitatory equations, Δt is the time of post-synaptic neuron firing minus the time of presynaptic neuron firing. There is only one *A* variable for inhibitory edges, as potentiation and depression are weighted equally (i.e. they have equal amplitudes). Similarly, there is only one *τ* time constant, as the STDP curve is symmetric along the y-axis. *η* is a variable unique to inhibitory edges and it controls the learning rate of the edges by scaling each weight change. *α* is a small value that is taken away from the weight each inhibitory edge when there is a presynaptic firing. This allows for Δw to be negative when Δt has a large magnitude in either direction. Also, for clarification, a negative Δw would result in a weaker inhibitory edge.

The learning parameters for our STDP rules were taken from data fit to physiological murine neuron experiments(50). Based on our STDP rule for excitatory edges, the chosen parameters reflected the amplitude of depression changes being greater than the amplitude of potentiation changes(1, 51). This allowed for depression to control the activity of the network to avoid signals propagating indefinitely. For the STDP rule for inhibitory edges, the chosen parameters weighted potentiation and depression equally, as there was no reason to augment inhibitory edges differently based on whether they were successful at reducing network activity or not(52).

The geometric embedding of our model is quite consequential here, as each edge having a specific length varies the amount of time it takes for signals to reach their target neurons(53). All signals travel at the same speed, so the offset in their arrival is based solely on these lengths. The delays along the edges are taken from a normal distribution around 730*µ*m(44) with a conduction velocity of 1.68 m/s(45).

The weights of the edges in the network are chosen from a uniform distribution, such that 80% of the edges are excitatory and range from the threshold potential down to just above 0 and 20% of the edges are inhibitory and range from the negative magnitude of the threshold up to just below 0. This results in single neurons having both inhibitory and excitatory outgoing edges, rather than being explicit inhibitory or excitatory neurons themselves. We decided to make 20% of the edges (synapses) inhibitory, as that has been observed in the rat hippocampus(46). Neurons do not normally have both excitatory and inhibitory outgoing edges due to Dale’s law, so we assumed inhibitory edges contained length-less inhibitory interneurons to simplify the model(54). These inhibitory interneurons needed axon lengths of 0 to keep the average delay of inhibitory edges the same as the average delay of excitatory edges. Excitatory and inhibitory edges used the same ranges for their weights, as this equalization has been observed in murine pyramidal neurons(55).

To improve computational efficiency, we use an event-driven simulator instead of integrating the neurons over time(56). With this, computations are done every time a signal arrives at a neuron. This allows for refractory time, potential summation, and membrane leak to be calculated only when necessary.

### B. Network Model

We use a graph abstraction to describe the structure of the network. A directed graph *G* consists of vertices *V* which may be connected by directed edges *E*. The dynamics on the graph follow that of a biological neural network, where each vertex represents a neuron and each edge represents an axon. We abstract a couple of salient features of the biological neural network such as edge delays and edge weights and directly encode them in the structure of our network. Each edge is endowed with a delay *d*_*ij*_ whose subscripts *i* and *j* represent the initial and terminal vertices, respectively. The edge delay represents the duration of the traversal of an action potential along the axon of a neuron. The value of the edge delay accounts for the conduction velocity of the action potential and the physical distance the action potentials must traverse between neurons. Each edge is also endowed with a weight *w*_*ij*_, which represents the synaptic weight and is modified by the rules stated by the neuron model. In Section A, we describe in detail how we ascertained the values for *d*_*ij*_ and *w*_*ij*_.

Our network consists of two distinct layers of vertices; we call them the input layer and the recurrent layer. We do not have an output layer as most other approaches do. Each layer is distinguished by its connectivity, number of vertices, and initial values. Vertices of the input layer have edges connected in a feed-forward manner, such that each input vertex is connected to all recurrent layer vertices. Each vertex of the recurrent layer is connected to every other recurrent layer vertex, which results in a complete subgraph *G*^*hid*^ ⊂ *G*.

### C. Inputs

To stimulate our network, we use images from the MNIST data set from the National Institute of Standards and Technology(57). This data set contains images of handwritten digits between 0 and 9. Each image consists of 784 pixels in a 28 by 28 grid. We map each pixel to one node within the input layer of our network and every non-zero pixel causes the corresponding input node to fire when the simulation starts. As such, pixels with the maximal intensity of 255 are treated the same as pixels with an intensity of 1. All appropriate input nodes fire instantaneously at time 0, but their signals reach neurons within the recurrent layer at different times depending on each edge’s distance and conduction velocity. The network is only stimulated one time by this process and then the dynamics within it are allowed to run undisturbed.

**Table 1.** The parameters used within the network.

|  |  |
| --- | --- |
| Refractory Period(42) | 3.9 ms |
| Resting Potential | 0.0 |
| Threshold Potential | 2.0 |
| Upper Bound for Edge Weight | 2.0 |
| Lower Bound for Edge Weight | -2.0 |
| Mean of Edge Length Gaussian(44) | 730μm |
| Standard Deviation of Edge Length Gaussian | 219μm |
| Speed of Signals Across Edges(45) | 1.68 m/s |
| Membrane Potential Leak Time Constant(50) | 0.0093 |
| Excitatory STDP Potentiation Amplitude(50) | 0.00008 |
| Excitatory STDP Potentiation Time Constant(50) | 7 |
| Excitatory STDP Depression Amplitude(50) | 0.00014 |
| Excitatory STDP Depression Time Constant(50) | 10 |
| Inhibitory STDP Potentiation Amplitude(47) | 0.2 |
| Inhibitory STDP Potentiation Time Constant(47) | 20 |
| Inhibitory STDP Depression Amplitude(47) | 0.2 |
| Inhibitory STDP Depression Time Constant(47) | 20 |

Each of the parameters chosen for the latencies in the network and the STDP rules was taken from murine hippocampus experimental data. The only exception is the edge weight range and threshold potential, which were chosen to make the computations manageable.

### D. Simulation Observables

We analyze the state-space trajectory of edge weights to determine whether images from the same class result in similar dynamics. Initializing the network with a sample MNIST image causes a subset of the input layer vertices to activate. When a vertex activates, it generates a signal on each of its outgoing edges. These signals either cause downstream vertices to activate or have no effect if the receiving vertex is refractory. Since every input layer vertex is connected to every recurrent layer vertex, there will be very few cases where the signals arrive, but then simply decay away completely over time. In either case, the edge synaptic weight is modified by the rules of STDP described in section A. The changes in synaptic weight encode the dynamics on the network induced by the stimulus.

The vector of synaptic weights can be written as 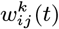, where *ij* identifies the edge for the *k*^*th*^ MNIST simulation. To calculate the classification accuracy, we sample the vector of synaptic weights every 100*ms*. We do not consider the synaptic weights of the edges between the input layer and the recurrent layer, but only the synaptic weights of the edges within the recurrent layer itself. This is because we are focusing on synaptic changes caused by activations within the recurrent layer, rather than directly from the image itself. As the input is only fed in once, there would only be one STDP change at most per edge between the input and recurrent layer. Therefore, for *n* recurrent layer vertices, the size of the synaptic weight vector at each sampling time is *n*(*n* − 1).

For the classification task, we compare the vectors of synaptic weights at each sampling time. The feature vectors of *K* MNIST simulations can be written in the following matrix notation:

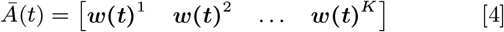

*Ā* (*t*) tracks the evolution of the feature vector over time.

### E. Parallel Embedding of Inputs

Each input to the network is treated independently. When we present the network with an image, each input layer neuron is activated only once when its corresponding pixel has a value *>* 0. We do not present periodic, constant, or random rate-based inputs to the network. Upon stimulation with an image, we record the dynamics over the course of 2*s*. From each simulation, we analyze the state-space trajectory of edge weights. Since each simulation is independent of the others, this allows us to simulate many networks in parallel. As such, they are not sequentially trained. Each parallel simulation begins with the same initial conditions. The number of parallel simulations is only constrained by computational resources.

We analyzed 10, 000 MNIST images in this manner and compared vectors of synaptic weights between them. With only one stimulation at the appropriate input neurons, the simulations themselves simplify down to volleys of activity within the recurrent layer. Instead of training individual networks on specific classes, we compare the resultant synaptic weight vectors to determine which class they are most similar to. This, therefore, informs which digit the input must have been.

### F. KNN for Classification Accuracy

Once we obtain the vectors of the synaptic weights from the many simulations, we run them through a weighted k-nearest neighbors (w-KNN) algorithm to determine their localities(58). Using w-KNN provides us with a simple way to assess synaptic weight vectors based on their Euclidean distances to discern if they are of the same class or not. In this w-KNN, we classify each vector in the test set based on their distances from the 9, 000 vectors in the training set. The w-KNN weighting of each neighbor comes from the inverse of the distance between it and the weight vector being evaluated; whichever MNIST digit has the highest total weight based on the 5-nearest neighbors is the one that is chosen for the classification. The 10, 000 total vectors are sorted into the training and test sets randomly and used for classification over 10 iterations to avoid any sampling bias. In addition, because there are approximately the same amount of simulations per digit in our population of 10, 000 vectors, imbalances should not create biases in modes when many neighbors are considered. Importantly, even though the neighbors are pulled from a “training set”, there is no actual training in this system, as all of the simulations were performed independently.

### G. UMAP for Visualization

In order to visualize the clustering of these synaptic weight vectors, we use the UMAP dimension reduction algorithm (59). We take edge weight vectors from 10, 000 simulations and run them through UMAP to see how separated each digit becomes from the others. UMAP employs k-nearest neighbors concepts with a unique local metric space implementation to effectively reduce the dimensionality of large data. By using this, we obtain a two-dimensional plot of all of the vectors labeled by their input digit. This reveals to us how similar or distinct the state-spaces of vectors of each digit can be based on their proximity to each other.

## 2. Results

### A. Characterization of Network Activity and Embedding Quality

When implementing our model, our first focus was on characterizing the activity within the network. To accomplish this, we recorded the time of the first 30 activations for each recurrent layer neuron in a 200-neuron network. As shown in Figure 3, the activity is not identical across all neurons, but we can see that most neurons have fired 30 times by 300 ms post-stimulation. As shown in Figure 4, information seems to be embedded best at 300 ms post-stimulation. That means the network is not just embedding a few key activations that reflect the stimulus directly, but rather aggregating thousands of activations into the structure of the network. Additionally, each activation leads to hundreds of synaptic weight changes via the STDP rules.

**Fig. 3.**
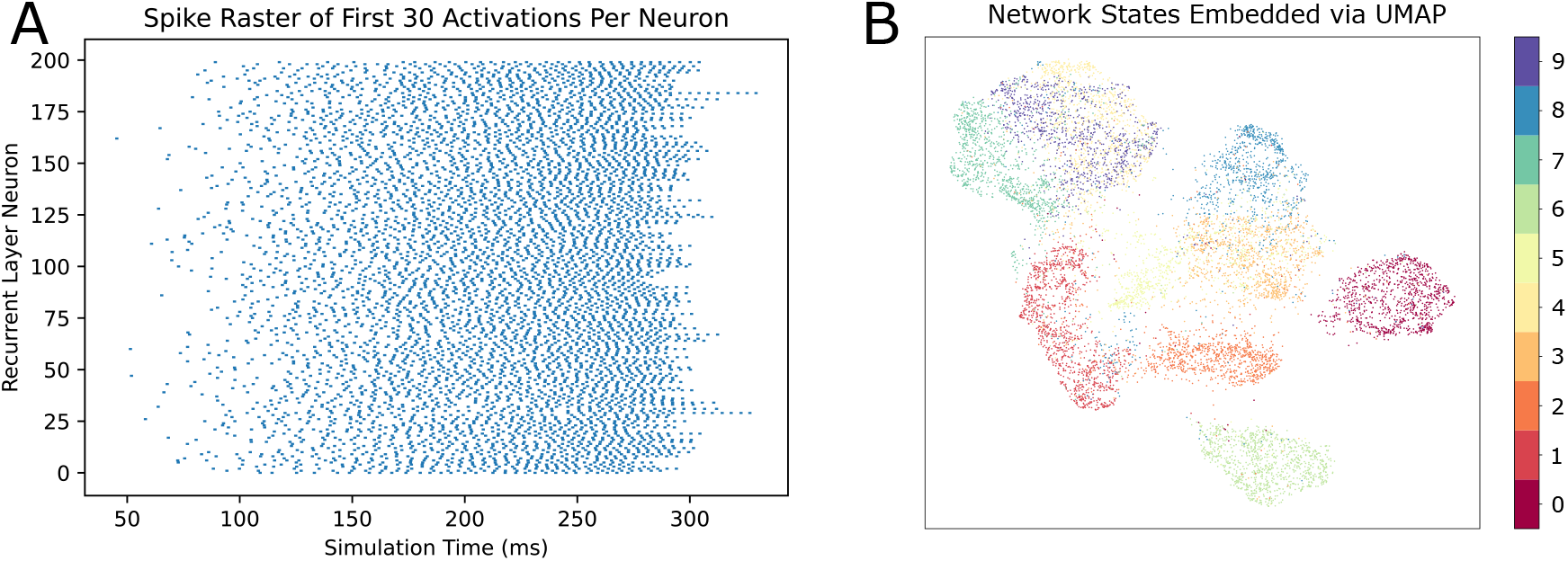
(A) It can be seen here that the activity of the network starts slowly due to the delays from the input nodes, but quickly increases. In addition, even with just 200 recurrent nodes, there seem to be as many as 20,000 activations per second in the simulation. As such, the activation dynamics are constantly changing the weights of the system. (B) Using a UMAP projection to reduce the dimensionality of the edge weight vectors, we can get a glimpse of how they may be separating from each other over the course of the simulations. Edge weight vectors from stimuli of the same digit generally congregate together in this projection, demonstrating that they may be more similar to each other than to vectors from other digits.

**Fig. 4.**
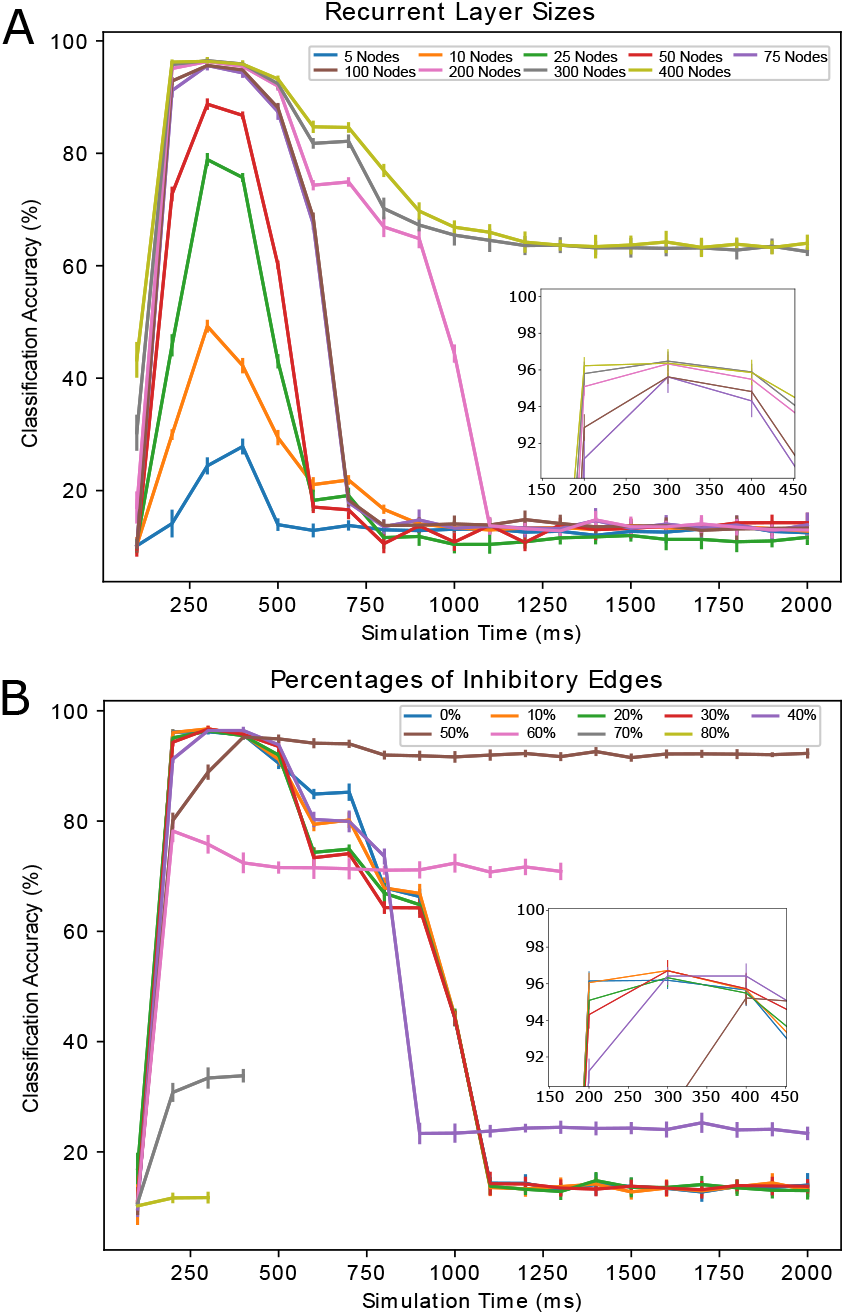
(A) We tested various recurrent layer sizes to balance the classification accuracy with simulation efficiency. The peak classification accuracy improved as the size of the network grew, but remained at 300 ms of simulation time. However, we observed that the improvements gained in networks larger than 200 nodes were not substantial at peak classification accuracy, but rather when the accuracy began to diminish. (B) We tested different percentages of inhibitory edges within the network. Some percentages, such as 70% and higher, were so negative that they silenced the network prematurely. When comparing the peak classification accuracies across all percentages, we observe that 20% has the highest peak. Interestingly, some inhibitory percentages have peak accuracies at simulation times later than 300 ms, which is a phenomenon that is not observed with many other parameter configurations.

To get an idea of the behavior of networks stimulated by different images of the same class, we took edge weight vectors at different simulation times and ran them through a UMAP dimension reduction. The greatest separation occurred at 300 ms post-simulation, as seen in Figure 3. Notably, even though the networks were stimulated independently, they tend to congregate together based on the class of the image used. To generate Figure 3, only the edge weight vectors were used with no additional information about spiking activity or stimulus pattern. Within the structural embedding itself, there was information about the stimulus. Further, we did not include edge weights between the input and recurrent layer, as they would not reflect the activity within the recurrent layer. Instead, signals would only travel along those edges due to image pixels and the edges, therefore, would only be affected by STDP once. As such, these STDP changes would be direct reflections of pixels from the image and would not demonstrate any embedding due to network dynamics. As we are interested in determining how well network dynamics can embed information through thousands of signals involved in STDP, these input layer to recurrent layer edges would not contribute overall.

Based on the UMAP data supporting our hypothesis that the edge weight vectors contained information themselves, we used a k-nearest neighbors algorithm to try to classify networks with unknown stimuli based on their edge weight neighbors. As the information capacity of the edge weight vector is bound to its dimension, we decided to experiment with the number of neurons (and therefore, edges) within the recurrent layer. As shown in Figure 4, as the network size grows and the number of edges increases, the peak classification accuracy of the networks using a k-nearest neighbors algorithm increases. With 200 neurons in the recurrent layer, the peak classification accuracy was 96.5±0.5%. With networks larger than that, peak classification marginally increased while substantially enlarging the edge weight vector space. With *n* recurrent layer neurons, an edge weight vector would contain *n*(*n* − 1) edges. Though a recurrent layer of 300 neurons has slightly higher accuracy, the computational costs are disproportionately larger. For all network sizes, the peak classification accuracy was reached at 300 ms post-stimulation, which is an interesting phenomenon independent of the number of neurons. As the network size increases, the embedding of the stimulus seems to be retained longer within the network. With recurrent layers as large as 300 or 400 neurons (89,700 or 159,600 edges), the fall in accuracy did not even occur within the 2-second window we were using.

To continue our characterization of this recurrent layer, we also changed different percentages of the edges to be inhibitory. For a network with a 200-neuron recurrent layer, the peak classification accuracy of 96.5±0.5% was achieved with edge inhibitory percentages of 10-30%, as can be seen in Figure 4. This matches biological observations of inhibition quite nicely(46, 60–62). Interestingly, with 50% of the edges being inhibitory, the network neither becomes quiescent nor loses the embedding of the stimulus over time. While it does not achieve the same peak classification accuracy as lower inhibitory percentages, this maintained embedding could be useful in some biological systems.

### B. Extending Simulations Beyond Experimental Plausibility

To properly evaluate the contribution of both potentiation and depression to the network embeddings, we then ran simulations with each separately for comparison. As shown in Figure 5, turning off potentiation within our networks actually had a minimal effect on the peak classification accuracy. With just depression gradually decreasing the edge weights, the network could still embed the stimulus with the same success. However, once we turned off depression, potentiation alone was incapable of maintaining the embedding quality of the stimulus. With depression and with or without potentiation, a classification accuracy of 96.36±0.47% was achieved with a recurrent layer of 200 neurons. In contrast, without depression, only a classification accuracy of 87.24±0.78% was achieved with the same sized network.

**Fig. 5.**
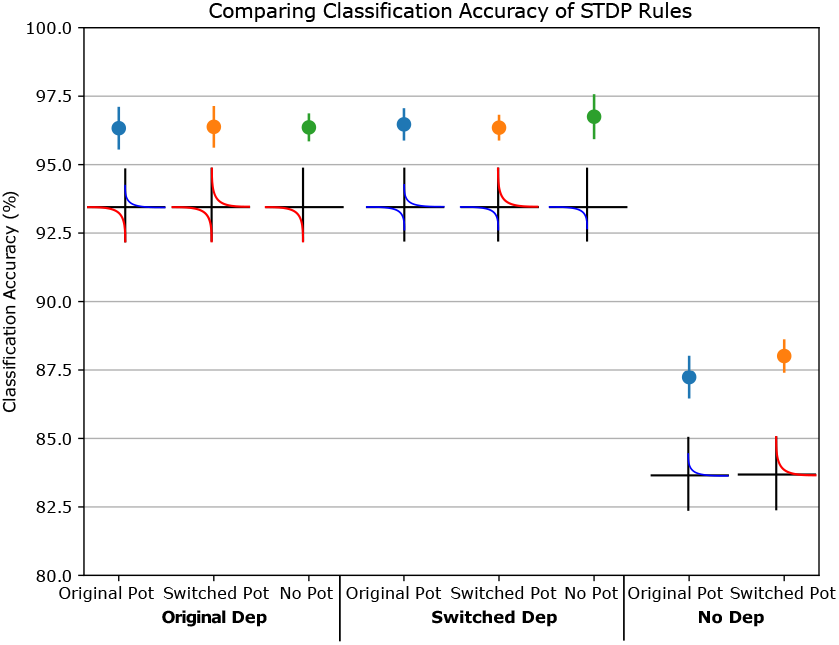
We tested the importance of potentiation and depression on the edge weight changes. By turning each of them off separately, we see that just depression is sufficient to maintain high classification accuracy, but just potentiation reduces the peak accuracy substantially. We then switched the parameter values for each learning rule to see if they were the cause, but we observed the same phenomena regardless of which parameter values were used for each learning rule. As such, edge weight depression seems to be necessary for the stimulus embedding in this system.

To disqualify the effect of the parameter values themselves (which were observed experimentally), we switched the parameters used for depression and potentiation as well. Originally, the parameters had depression with changes of higher amplitude than potentiation. As Figure 5 shows, even after this was switched, depression still had a larger impact on classification accuracy. In any paradigm without depression, the accuracy would drop from 96% to 88%. On top of that, paradigms without potentiation maintained 96% accuracy as long as depression was working.

To further investigate the balance between potentiation and depression, we next changed the refractory period of the neurons in the recurrent layer. As the frequency of potentiation/depression events depends on the length of the refractory periods, it follows that this would have a large impact on which piece has more influence. Interestingly, for all the refractory periods we tested, from 100 ps to 10 ms, having both potentiation and depression changing edge weights maintained maximum classification accuracy in our system as seen in Figure 6. It is only when looking at potentiation or depression alone that differences became noticeable. With shorter refractory periods, just potentiating changes performed better than just depressive changes. However, once refractory periods grew longer and more depressive events took place, only depression performed better than only potentiation. Across all tested refractory periods, having just one or the other was sufficient to reach the maximum classification accuracy obtained by having them both together. In addition, the inflection point where depression becomes more impactful than potentiation seems to be with a refractory period of around 1 ms.

**Fig. 6.**
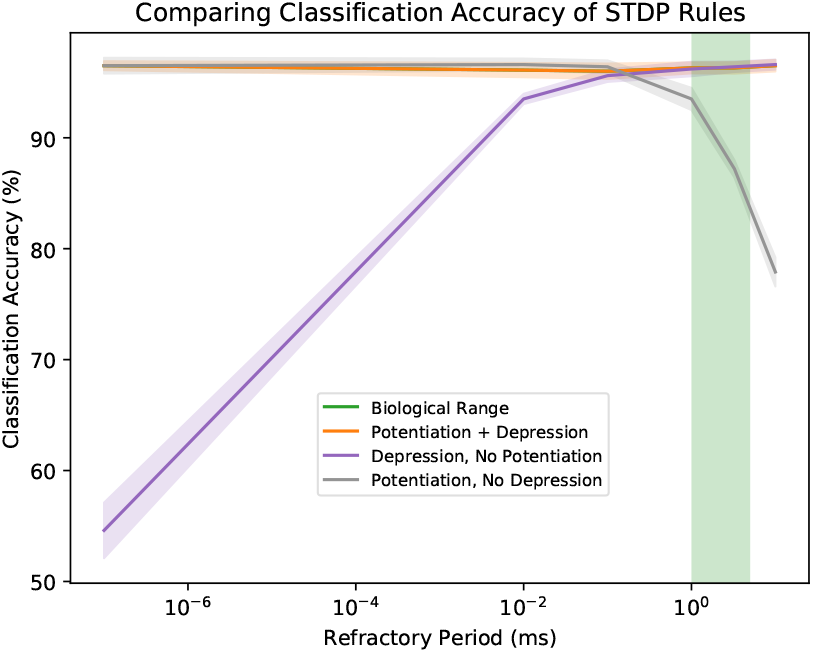
We increased and decreased the refractory periods of neurons within the recurrent layer to observe if there was a quantifiable reason for the scale of biological refractory periods. Interestingly, the first thing we see is that regardless of how large or small the refractory period is, with both potentiation and depression changing weights, the classification accuracy stays remarkably consistent. However, with just depression affecting the weights, the longer refractory periods allow for better classification accuracy. Inversely, with just potentiation, the shorter refractory periods lead to better classification accuracy. There seems to be an inflection point at around 1 ms where depression alone begins to embed information better than potentiation alone.

### C. Comparing Edge Weight Vectors to Image Stimuli

As we studied the nearest neighbors for each simulation and how they seemed to well-represent similar stimuli, we decided to look more closely at the raw images. As shown in Figure 7, we plotted the 9000 nearest neighbors from one particular MNIST image based on pixel value distance. Then, over the course of a simulation, we plotted the 9000 nearest neighbors of that same image based on the edge weight vector distance. Taking the running mean of each group of 100 edge weight vector distances, we can see similar patterns start to form and disappear over the course of a typical simulation. Notably, the edge weight vector distances seem to correlate with the pixel value distances between 200 and 400 ms of simulation, when peak classification accuracy occurs. After 400 ms, the edge weight vector distances lose correlation with the pixel value distances.

**Fig. 7.**
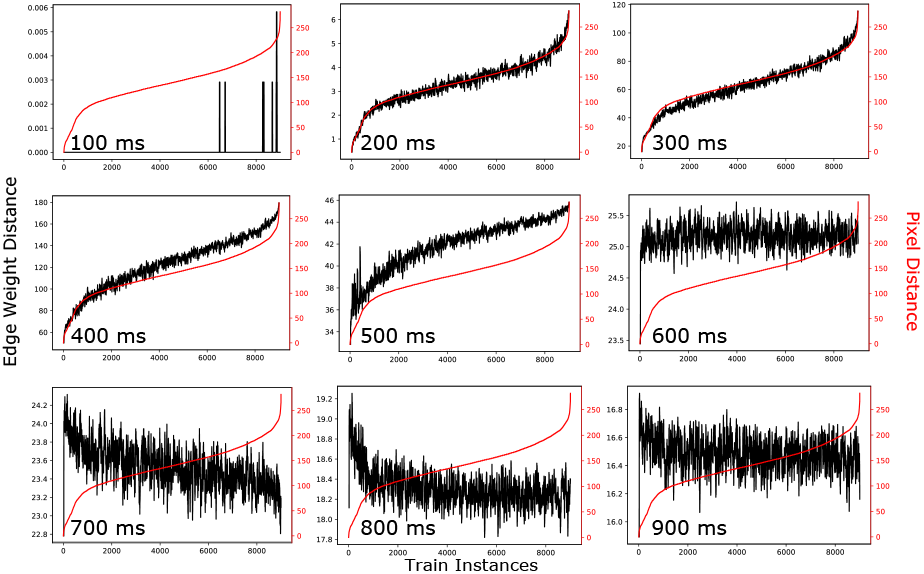
In red, we see the distances of the neighbors to a single image based on their raw pixel values sorted to be ascending. Over the course of the simulation, we use this ordering to see how it compares to the distances to the same image based on the network edge weights (in blue) while preserving the order from the raw pixel values. The running mean of the nearest 100 image edge weight distances is used to produce the orange line. We can see that around 300 ms, where the embedding is at its best, the orange edge weight distances seem to correlate nicely with the red raw pixel distances. However, this correlation is transient and is not reached before 200 ms of simulation time nor preserved after 400 ms.

## 3. Discussion

### A. Parsimony Between Our Results and the Neurobiology

Broadly, our results coincide with the expectations defined by neuroscience while also providing insights in a way only computational models can. The parameters taken from the neurobiology allowed us to employ plausible learning rules and time delays while the KNN algorithm allowed us to view information embedding capabilities in ways that cellular experimentation may not be able to currently. Through this, we aligned our results with widely accepted assumptions, such as networks needing to be sufficiently large to embed a stimulus(38) and inhibition playing an essential role in information embedding(4). On top of that, we demonstrated the effectiveness of structural embeddings in networks using STDP rules. Our work extends the idea of structural changes being a part of the process of embedding memories(8) to them being possibly sufficient to do so on their own.

As shown in Figure 3, our networks are full of activity. Therefore, the embeddings we observe are compositions of large combinations of weight changes. Rather than a few highly specific activations triggered by the stimulus, thousands of activations are integrated to produce the network state that is used. Without any pre-training, the base network stimulated independently with two images of the same class will likely end up with two similar network states 300 ms into the simulation. Further, images with various orientations across the same class share similar enough patterns to trigger similar weight changes overall. Figure 3B demonstrates this, as states of networks of the same classes aggregate at particular times when simulated independently of each other. It is possible that some networks in the brain may work similarly, particularly in the perirhinal cortex, where object recognition is accomplished by networks that have strong depression, as our networks do(22). In the brain, similar stimuli may result in similar structural states for object identification.

As shown in Figure 4A, our simulations demonstrate the notion that networks that are too small may not be capable of reconstituting a stimulus fully(38). However, our simulations explore the trade-off between stimulus embedding and network size. As we increase the size of our networks beyond 200 recurrent neurons, the classification accuracy improves only marginally while increasing the computation time dramatically. When thinking about the tangible resources in the brain, such as neurotransmitters and ATP, it could be inefficient to use larger networks for problems, as the embedding capacity may improve marginally at an enormous cost. Others have observed this phenomenon in working memory; training increases the number of neurons initially stimulated, but it does not increase the number of neurons recruited to the assembly in general(40). While raising the number of neurons that are initially stimulated increases the ATP consumption, this increase is less than that by enlarging the assembly itself. Therefore, there is a more efficient trade-off between increasing resource usage and computational capacity in these neuron assemblies.

The importance of the right amount of inhibition was also echoed in our simulations. While experimental observations have highlighted key inhibitory percentages that are often observed in the brain(46, 60–62), our network allowed us to take that to its extreme as well. We noticed the best classification accuracy with inhibition percentages between 10-30%, which aligns with what is seen experimentally. Our observations support our hypothesis that the brain may have selectively evolved networks with these inhibitory percentages to embed information optimally. However, we also observed a strange phenomenon with 50% inhibition that is not easily explained. The activity of these networks did not diminish due to inhibition, nor did their classification accuracy decrease significantly after hundreds of milliseconds of simulation. One possible function for a network with this inhibitory percentage is the acquisition and maintenance of fear memory through inhibitory neurons in the intercalated cell masses (ITCs) of the amygdala(63). These masses are densely inhibitory and connect the amygdala to surrounding structures. In our model, all inhibitory neurons follow the same activation and plasticity rules rather than considering their true variability. *In vivo* experiments demonstrate different populations of inhibitory interneurons working independently of each other (64, 65). Perhaps the phenomena we observe with 50% inhibition resulted from an oversimplification of the different populations in biological inhibition.

### B. Extending Our Model to Better Understand Biological Networks

Since our simulations allowed us to manipulate neuron refractory periods, we explored the reasoning behind the biological refractory period. While neurons cannot employ shorter refractory periods due to the kinetics of sodium and potassium channels, it does not seem like it would be disadvantageous to do so if it was possible(20). Based on our simulations, the benefits of an absolute refractory period of 1 ms are not immediately apparent. While a refractory period of that duration allows for depression to have a more prominent role, with shorter refractory periods, potentiation could take over without losing information embedding capability. However, too short of a refractory period could lead to hyperactivity of networks, which would require excess resources without a clear improvement in embedding capability. Inversely, longer refractory periods would also preserve embedding capability but may cause neurons to be too slow to respond to new stimuli. A previous experimental and computational study on retinal ganglion cells noted the benefit of a sufficiently long refractory period to allow for reproducible responses within the network(66). It is possible that outside of the ion channel limitations, the biological refractory period was optimized for reproducibility rather than embedding capability. Further, research has been done on the importance of the relative refractory period when modeling cortical data(67), which may also be an important consideration when interrogating these

#### STDP dynamics

In addition to changing refractory periods, our simulations allow for the isolation of potentiation and depression, which is experimentally challenging to do *in vivo*, to demonstrate the importance of depression. In Figure 6, we demonstrated that with a biological refractory period, depression alone embeds a stimulus better than potentiation alone. Similarly, in a mouse model, it has also been observed that potentiation alone is less effective at embedding working memory than when both potentiation and depression are active together(68). Based on the different refractory periods we tested in Figure 6, we see that potentiation and depression alone consistently embed information at different efficiencies. With shorter refractory periods, potentiation works better alone than depression while with longer refractory periods, depression works better alone than potentiation. This observation is intuitive, as fewer signals arrive at refractory postsynaptic neurons with a shorter refractory period and, therefore, those signals could contribute to future activations. Another consideration for an optimal refractory period is the relative strengths of potentiation and depression. Experimental data has shown depression to have a larger magnitude of change than potentiation(1, 3, 69). However, this may just be a consequence of the refractory period, as our data shows that depression has stronger embedding capabilities than potentiation with a biological refractory period regardless of plasticity parameter values. As such, the magnitudes of depressive changes may seem larger experimentally because the embedding capability of depression is better at that refractory period. Quantitative experimental observations may be influenced by the efficacy of depression at embedding information with a biological refractory period. In addition, since our model shows the importance of depressive events for embedding, we speculate that the biological magnitude is larger for depression than potentiation to increase the fidelity and range of the depressive weight changes. This is a beneficial adjustment as it could allow for embedding through unique, subtle depressive changes to weights. In a hypothetical brain consisting of neurons with shorter refractory periods, the magnitudes of change for potentiation and depression may appear different, as the mechanisms behind them are time-sensitive. It is possible that a refractory period of 1 ms allowed for the optimal balance between the aforementioned reproducibility and the magnitudes of potentiation and depression.

The analysis of the synaptic weight space of our networks aligns well with studies into the state spaces of memory(70, 71). It has been proposed that networks in the brain may alternate between persistent states to store memories. These states could possibly consist of synaptic weights, synaptic connections, membrane voltages, neuronal activation patterns, or other biochemical variables. In our model, we directly use synaptic weights to define the different states that networks transition between as they are stimulated. In particular, we show that networks stimulated by images of the same class reach similar states after 300 ms of simulation, as seen in Figure 3. One natural question that arises from the synaptic weight state space is its potential capacity. We show that there is a useful amount of separation with ten classes, but further work could be done to see if there is separation when the number of potential classes increases. In addition, these synaptic weight states do not seem to persist over time, so the dynamics that lead to its transience may also be further studied. Perhaps if there was a glial component to the network, the states would be supported to persist for longer. Experimental studies have shown that glial cells play a key (though not yet well understood) role in hippocampal structural maintenance(72, 73). Incorporating glial cell support could allow the weight changes in our network to preserve information for longer and delay our observed loss in accuracy.

In addition to support, glial cells may function as downstream readers of synaptic weights in the hippocampus. These cells could play a role reminiscent of our KNN algorithm. In particular, astrocytes extend processes to synapses within the hippocampus to monitor and contribute to the activity and plasticity that takes place there(74). While the astrocytes may not keep track of the absolute synaptic strengths themselves, their internal calcium gradients change with potentiation and depression of the synapses(48, 75). As such, their internal states integrate the overall plasticity changes over thousands of synapses to monitor the population as a whole(13). As astrocytes can communicate with each other through gap junctions, they can transfer this information across the syncytium to other astrocytes(14, 15). Those astrocytes, as a result, can further transfer that information to downstream networks of neurons by impacting their activity and plasticity(76). Astrocytes, therefore, can read the overall synaptic plasticity changes of a network caused by a stimulus (and possibly embedding that stimulus) and transfer it to another downstream network to facilitate the flow of information. Our KNN algorithm is similar, as it is sensitive to the edge weight changes within the networks rather than their absolute weights, which are largely influenced by the initial state of the network. The synaptic plasticity information astrocytes integrate could be the biological analogue of our edge weight vectors and could serve as a plausible mechanism through which information is transferred in the hippocampus.

Furthermore, this view of astrocytes as readers of synaptic weights may explain the cause of transient epileptic amnesia(77). When considering the paradigm of rate coding, seizures represent a unique firing rate for various regions of the brain. Despite that, most patients still suffer amnesia during the episode and often in moments subsequent to it(78). This would imply that there is more to memory formation than firing rates alone, as epileptic firing rates would not be confounded by any other event. This is where our hypothesis fits in nicely. The chaotic network activity of seizures may cause STDP changes that do not properly embed any particular stimulus. As such, when the brain tries to recall memories from the event, it may not be able to accurately process what the synaptic changes represent. Morphological changes have been observed in the synapses of epileptic brain regions(79), so there is precedent that seizures abnormally change synaptic plasticity. In addition, there may be dysfunction in the level of astrocytes as well. As astrocytes are critically important for removing excess neurotransmitter from synapses, seizures due to insufficient removal may be caused by astrocyte dysfunction(80). This is uncontroversial, but if we view astrocytes as readers of synapses as well, we may understand transient epileptic amnesia better. If issues arise in astrocytes maintaining synapses, issues may also arise in astrocytes being able to read synapses. This could hinder the transfer of information across networks in the brain and would, therefore, impede both memory formation and recall. While there have been hypotheses for how amnesia arises with rate coding, such as misaligned gamma and theta wave synchronization(79), viewing astrocytes as key players in storing memory elucidates one more possibility as to why memory issues persist.

### C. Future Directions For This Work

In the future, the introduction of noise into the system could be studied as well, as experimental studies have shown STDP to struggle when noise via jitter is added to the spike timing(81). Inversely, adding minimal noise may be beneficial to allow subthreshold signals to impact network dynamics(82). Further, our threshold potential was chosen such that an average of three excitatory signals would be required to reach it. Follow-up experiments could adjust the threshold potential to observe how increasing and decreasing the number of activations affects the STDP embedding of the stimulus. Lastly, another future direction could involve studying the effect of multiple sequential stimuli on the embedding. If the synaptic weights are capable of reflecting one stimulus, what happens when two are presented? Do the weights reflect a composition of the inputs?

When looking at neurons within the brain, our synaptic weight spaces offer valuable insight. While rate coding, postsynaptic potential magnitudes, and myelin thicknesses are primarily considered as evidence of learning(5, 6, 16), structural changes, in particular, can be meaningful as well. Our methodology demonstrates that the stimulus can be identified by interrogating the structural state of a network following stimulation. In addition to astrocytes potentially monitoring synapses, the structural state may also strongly influence the overall rate and magnitudes of activations within the network. As such, the structural state may correlate well with current observations of activation rates and postsynaptic potential magnitudes. As brain slice imaging techniques continue to improve, subtle structural differences associated with different learned patterns may also become observable experimentally. When designing our network, we used a single recurrent layer to avoid any brain region-specific architecture. To generalize functions and interactions of various brain regions, we minimized topological variability by connecting each input neuron to every recurrent layer neuron. However, various regions of the hippocampus and parahippocampal cortex are recurrent networks, so our model could represent an apt analogue for them(83, 84). In addition, we maintained the same initial conditions for simulations run in parallel (axon lengths, starting weights) to minimize variability. To properly evaluate our edge weight and delay distributions, we tested 11 randomly chosen initial conditions and observed a classification accuracy of 96.53±0.06% with a 200-neuron recurrent layer. With these controls in place, the experimental data we gathered resulted from changes to the refractory periods, network sizes, inhibitory percentages, and STDP rules. Regarding this design philosophy, further work can be done to investigate how different network architectures are impacted by varied refractory periods and potentiation/depression isolation. Furthermore, allowing for variability in initialization or even sequential stimulation could lead to new insights into the homeostatic capabilities of the networks to recalibrate and resume learning. To add more neuroscientific features to the network, we could also add dynamic dendritic spine calcium concentrations, as a recent model observed their effect on STDP strength(85). In particular, in high-activity networks such as ours, the magnitude of STDP changes could be quite variable depending on the calcium transients.

Experimental techniques capable of observing the graded breakdown of synaptic embeddings could produce diagnostic tools for neural dysfunctions and new understandings of computation in the brain. Research is currently being done to quantify how these synaptic embeddings relate to information storage(86), but it is still unclear how to best monitor them. In our model, embeddings are intimately tied to the timing of signals between neurons. Variability of a network’s geometry and signaling parameters affect timing, and volatility in either can cause failure in embedding robustness. Changes in a network’s geometry arise from available neural circuits due to disruptions to oscillatory neural patterns, and neuromodulatory signals (87–89). Dysfunction affecting the conduction velocity of axons, whether due to myelination or ion channel irregularities, could negatively impact memory storage due to inconsistent signaling parameters affecting the signal timings. Further, if there is limited neurotransmitter availability, either due to poor astrocyte recycling or insufficient production, the resulting variability in spike amplitudes will affect signal timing, thus the embedding, and may lead to memory problems. Another potential source of issues with memory could be connectivity changes due to synaptic pruning or neurotrophy, which could change the paths that signals travel along in response to the same stimulus. Lastly, conditions constraining the sizes of neuron assemblies could result in poor memory due to an insufficient number of edges on which information can be embedded. Biological experiments focused on any of these issues could elucidate mechanisms of memory loss beyond just synapse destruction due to neuroinflammation.

### D. Edge Weights As A Machine Learning Methodology

Stepping back from the biology, Figure 7 highlights a deeper dive into the correlation between the network embedding and the input images. In particular, after 300 ms of simulation time, the nearest neighbors to a specific vector closely resemble the nearest neighbors of the input image itself. This observation demonstrates the functional potential of our model. With STDP, neuron refraction, inhibitory edges, and edge latencies, the network recapitulates the stimulus after 300 ms.

This observation may imply that adding biological features to canonical artificial neural networks could benefit them for particular applications. However, this is not a novel idea, as many groups have already begun making neural network systems more biological with great success(90–93). Once our simulations have been completed, we use specific structural weight states from each simulation to classify. This reduces the number of parameters when compared to a deeper neural network(94).

In fact, since each simulation is run independently of the others, there is no “training phase” in our system. Each individual network embeds information in parallel via one-shot learning(94). The network activity of our model generates a dynamic embedding of the stimulus through the synaptic weight changes, which we have found to be empirically beneficial for classification. Our dynamic embedding can be viewed as a functional transformation of the network’s stimulus, where the function is stipulated by the individual network dynamics triggered by the stimulus. When we ran the KNN on the input images pixels we attained attained a classification accuracy of 94.5%, whereas, the KNN on our dynamic embedding resulted in a classification accuracy of 96.5%. Given that we empirically observed an increase in classification accuracy as a result of the dynamic embedding, we postulate that our transformation is non-trivial.

Further work can be done to investigate the machine learning potential of this system as well as other biologically-inspired systems. Although we only tested our network on the MNIST data set, which is no longer a hurdle for machine learning systems, more challenging data sets can be applied to it in the future. Classification accuracy of 96.5±0.5% is not as high as state-of-the-art neural networks achieve, but with one-shot learning, it is still a feat. In addition, due to the biological design, this network may prove advantageous for different challenges where conventional neural networks struggle.

## ACKNOWLEDGMENTS

We thank the Lawrence Livermore National Laboratory for providing us with extensive computational resources for our experiments.

